# A Novel Scavenging Tool for Cancer Biomarker Discovery based on the Blood-Circulating Nanoparticle Protein Corona

**DOI:** 10.1101/382192

**Authors:** Marilena Hadjidemetriou, Zahraa Al-ahmady, Maurizio Buggio, Joe Swift, Kostas Kostarelos

**Affiliations:** Nanomedicine Lab, Faculty of Biology, Medicine & Health, AV Hill Building, The University of Manchester, Manchester, M13 9PT, United Kingdom; Division of Cell Matrix Biology & Regenerative Medicine, Faculty of Biology, Medicine & Health, Biological, The University of Manchester, Manchester, M13 9PT, United Kingdom

**Keywords:** biomolecule corona, biomarkers, melanoma, lung cancer, nanomedicine ToC Graphic

## Abstract

The prominent discrepancy between the significant investment towards plasma biomarker discovery and the very low number of biomarkers currently in clinical use stresses the need for novel discovery technologies. The discovery of protein biomarkers present in human blood by proteomics is tremendously challenging, owing to the large dynamic concentration range of blood proteins. Here, we describe the use of blood-circulating lipid-based nanoparticles (NPs) as a scavenging tool to comprehensively analyse the blood circulation proteome. We aimed to exploit the spontaneous interaction of NPs with plasma proteins once injected in the bloodstream, known as ‘protein corona’ and to facilitate the discovery of previously unreported biomarker molecules for cancer diagnostics. We employed two different tumor models, a subcutaneous melanoma model (B16-F10) and human lung carcinoma xenograft model (A549) and comprehensively compared by mass spectrometry the *in vivo* protein coronas formed onto clinically used liposomes, intravenously administered in healthy and tumor-bearing mice. The results obtained demonstrated the ability of blood-circulating liposomes to surface-capture and amplify low molecular weight (MW) and low abundant tumor specific proteins (intracellular products of tissue leakage) that could not be detected by plasma analysis, performed in comparison. Most strikingly, the NP (liposomal) corona formed in the xenograft model was found to consist of murine host response proteins, as well as human proteins released from the inoculated and growing human cancer cells. This study offers direct evidence that the *in vivo* NP protein corona could be deemed as a valuable tool of the blood proteome in experimental disease models to allow the discovery of potential biomarkers.

**ToC Graphic:** 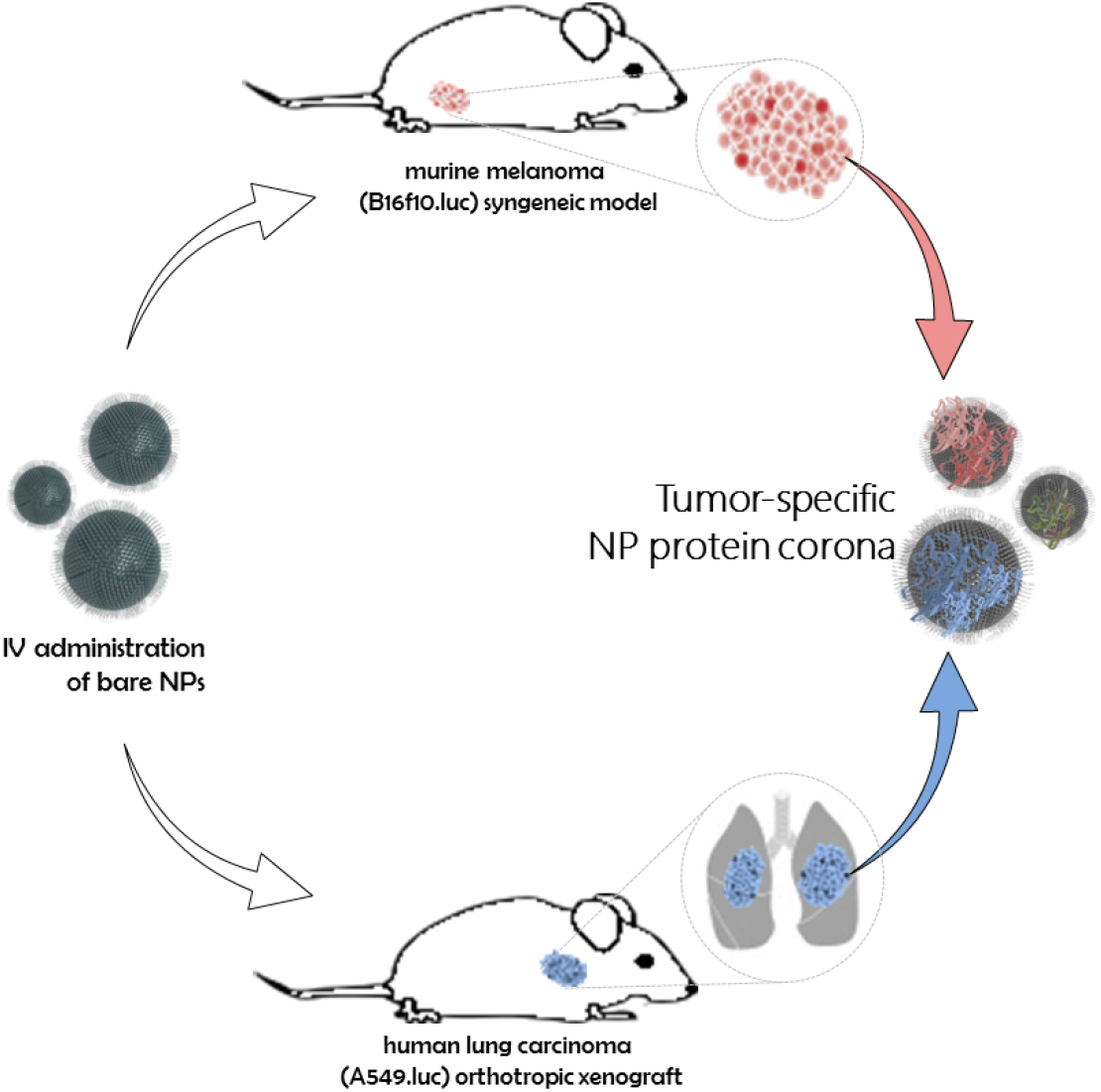

---

There is an urgent need to develop robust high-throughput proteomic platforms to enable the discovery of new molecular biomarkers for cancer diagnosis. Blood is considered to be the most valuable biofluid for protein biomarker discovery as it contains the most comprehensive human proteome, including also tissue leakage proteins that reflect ongoing disease states.^1^ However, discovery of biomarkers present in the blood by mass spectrometry-based proteomics has been shown to be particularly challenging and it is hampered by the large dynamic concentration range of blood proteins.^2^ Currently available proteomic tests, detect only a minute fraction of potential biomarkers due to their extremely low concentration in biofluids, in addition to the ‘swamping’ effect, caused by non-specific highly abundant molecules. The issue of signal-to-noise exceeds the current capability of proteomic analysis and therefore limits the diagnostic information that can be obtained.^3^

Nanotechnology-based platforms hold great promise in addressing the above fundamental and technical issues of biomarker discovery. Intravenously injected nanoparticles (NPs) could enable a thorough sampling of the blood proteome, considering their access to the host tissue microenvironments. The last decade, a vast literature has emerged aiming to examine the spontaneous interaction of NPs with plasma proteins once injected in the bloodstream, also named as ‘protein corona’.^4^ While the formation of protein corona was initially seen as a great hurdle that hampers the pharmacokinetic properties of nanomaterials, it is now seen as a tool with therapeutic and diagnostic potential.^5, 6^ Despite remarkable interest and a plethora of previous studies investigating the molecular composition of the protein corona formed around a variety of different NPs,^7-9^ to date there is no report in the literature demonstrating that intravenously injected NPs are able to surface-capture disease specific molecules.

In our previous reports,^10-13^ we developed a robust protocol to recover nanoparticles from the blood circulation of healthy rodents and investigated the previously elusive *in vivo* protein corona. *In vitro and in vivo* formed protein coronas were compared for the first time and the results demonstrated that *in vivo* protein corona was considerably more complex, consisting of many more different types of proteins in comparison to its counterpart *in vitro* corona.^10^ In a subsequent study,^11^ we employed doxorubicin-encapsulated clinically used liposomes to examine the time evolution of the *in vivo* protein corona. In addition to the highly dynamic nature of protein corona, the results demonstrated the tendency of liposomes to interact with a mixture complex of low abundance plasma proteins.^11^

Based on our previous findings, in the present study we hypothesized that intravenously injected liposomes are bound to low abundant disease specific proteins that cannot be detected by conventional proteomic analysis. We aimed to comprehensively compare the *in vivo* protein coronas formed onto intravenously injected liposomes in healthy and tumor-bearing mice and to investigate if corona fingerprints can be exploited for biomarker discovery in cancer diagnostics. PEGylated liposomes (HSPC:Chol:DSPE-PEG2000), that constitute the basis of the clinically used liposomal formulation Doxil^®^, were employed because of their established clinical profile.^14^ To prove our hypothesis, we employed two different tumor preclinical models: a syngeneic subcutaneous melanoma model (B16-F10) and human lung carcinoma xenograft model (A549). While syngeneic tumor models are representative of the host response in the presence of a functional immune system, inoculating human cancer cells into immune deficient mice creates a complex mixture of human cancer in murine stromal cells and potentially allows the simultaneous detection of tumor markers unambiguously secreted into the blood circulation from both non-cancer host stromal cells and human tumor cells.^15-17^

## Results

### Recovery of corona-coated liposomes from melanoma and lung adenocarcinoma-bearing mice

For the subcutaneous model, tumors were developed by injecting B16f10-luc mouse melanoma cells into the left leg of C57 female mice (n =3 mice/replicate; 3 independent replicates). For the lung carcinoma xenograft model, human A549-luc cells were intravenously injected into severe-compromised immunodeficient (SCID) male mice (n=3 mice/replicate; 3 biological replicates). After the transplantation of the two luciferase-expressing cell lines, tumor growth was monitored by whole body IVIS imaging (Figure 1A). When measurable bioluminescence signal was detected in all mice (2 weeks and 4 weeks post-injection of tumor cells for melanoma and lung carcinoma respectively), PEGylated liposomes (HSPC:Chol:DSPE-PEG2000) were intravenously injected (via the tail vein). Our previous time evolution studies in rodents proved that a complex *in vivo* protein corona was self-assembled around PEGylated liposomal doxorubicin (Caelyx®) as early as 10 min post intravenous administration. The results demonstrated that the total amount of corona proteins as well as the identity of corona proteins did not significantly fluctuate over time.^11^ For this reason in this study liposomes were recovered from the blood circulation of mice 10 min post-injection. Liposomes were also injected and recovered from healthy C57 and SCID mice as a control (n=3 mice/replicate; 3 independent biological replicates). Corona-coated liposomes were recovered from plasma and purified by unbound proteins by size exclusion chromatography followed by membrane ultrafiltration, as we have previously described.^10-12^

**Figure 1:**
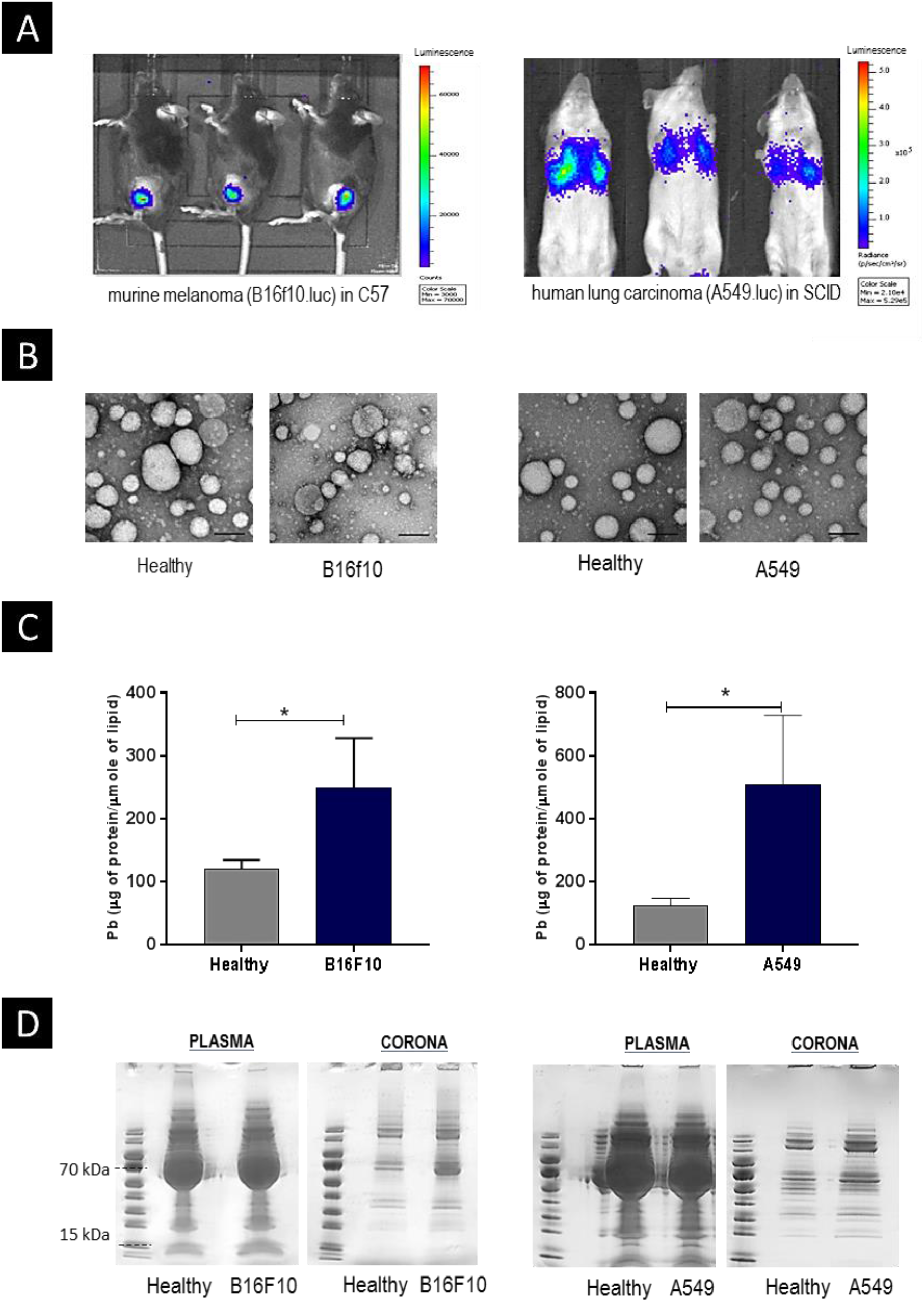
Protein Corona formation in tumor-bearing mice. **(A)** Representative bioluminescence images of melanoma-bearing C57 mice 2 weeks post subcutaneous injection of B16f10-luc cells and of lung carcinoma-bearing SCID mice 4 weeks post intravenous injection of A549-luc cells using the IVIS Lumina II camera; **(B)** Negative stain TEM of liposomes after their I.V injection and recovery from C57 and SCID mice. All scale bars are 100nm; **(C)** The total amount of proteins adsorbed *in vivo* onto liposomes recovered from the blood circulation of healthy and tumor-bearing mice. Pb values (μg of protein/μM lipid) represent the average and standard deviation from 3 biological replicates, each using 3 mice. *indicates *p<0.05;* **(D)** Imperial stained SDS-PAGE gel of plasma proteins control and corona proteins associated with liposomes.

The physicochemical characteristics of the PEGylated liposomes before and after their recovery from the blood circulation of healthy and tumor bearing mice are summarized in **Tables S1** and **S2**. A broader size distribution was observed by Dynamic Light Scattering of corona-coated liposomes in comparison to bare (non-coated) liposomes, whereas their surface charge remained negative (**Tables S1** and **S2**). In addition, Transmission Electron Microscopy showed a well-dispersed and protein-coated liposome population after recovery (Figure 1B).

The total amount of protein adsorbed onto the surface of liposomes was then quantified by BCA assay and Pb values were calculated (μg of protein/μmole of lipid). As shown in Figure1C, the average Pb values for tumor-bearing mice were higher than the Pb values observed for healthy mice, in both tumor models. These results provide initial evidence that protein corona fingerprints vary in the absence and presence of tumorigenesis. The higher amount of protein adsorbed onto liposomes in the highly vascularized lung carcinoma model in comparison to the melanoma model, further reinforced the hypothesis that protein corona formation depicts the ongoing pathophysiological aberrations reflected in the blood proteome.

Corona proteins associated with the surface of intravenously administered liposomes in healthy and tumor-bearing mice, were separated by SDS-PAGE, as illustrated in Figure 1D. In agreement with the quantification results, a higher total amount of protein adsorbed was observed in the presence of melanoma and lung-carcinoma tumors. The well distinct bands of corona proteins, even at the low MW region, demonstrated the ability of liposomes to minimise the noise of highly abundant proteins. Plasma samples from healthy and tumor-bearing mice were are also recovered by cardiac puncture as a control (without injection of NPs or any other processing). In contrast to the corona samples and as expected, the overwhelming signal of albumin in the case of plasma control was found to mask the low MW weight proteins (Figure 1D).

### Characterization of the *in vivo* protein corona in melanoma-bearing immunocompetent mice

The fact that albumin ‘masking’ was largely eliminated from the *in vivo* liposomal corona (Figure 1D) prompted us to investigate whether intravenously injected liposomes can form coronas with plasma proteins that cannot be directly detected by proteomic analysis of plasma control samples. Plasma control samples were prepared from blood collected from non-injected healthy and melanoma-bearing mice and without any further processing they were subjected to mass spectrometry analysis. To offer a comprehensive identification of proteins in plasma and corona samples LC-MS/MS was performed and data analysis was performed using two software tools, Scaffold and Progenesis. Scaffold software was initially used as a bioinformatics tool to identify and compare proteins in plasma and corona samples using relative protein abundance quantification, based on spectral counting. Processing of the raw data generated from LC-MS/MS with Progenesis discovery software was then carried out to more thoroughly, quantitatively and statistically, compare the levels of identified proteins in samples derived from healthy and tumor-bearing mice in to order discover potential biomarker proteins.

The Venn diagrams in Figure 2A show the number of common and unique proteins in healthy and melanoma-bearing C57 mice, as identified by mass spectrometry analysis of plasma control and corona samples. A significantly higher total number of proteins was detected in the corona samples (n= 932 in healthy mice; n=897 in melanoma mice) in comparison with the number of proteins identified by plasma analysis (n= 263 in healthy mice; n=225 in melanoma mice). In addition, the most abundant plasma proteins were not the predominant corona proteins as depicted in Figure 2B and **Table S3**. These findings demonstrate the ability of liposomes to capture and amplify low abundant blood proteins that cannot be directly detected by conventional plasma proteomic analysis.

**Figure 2:**
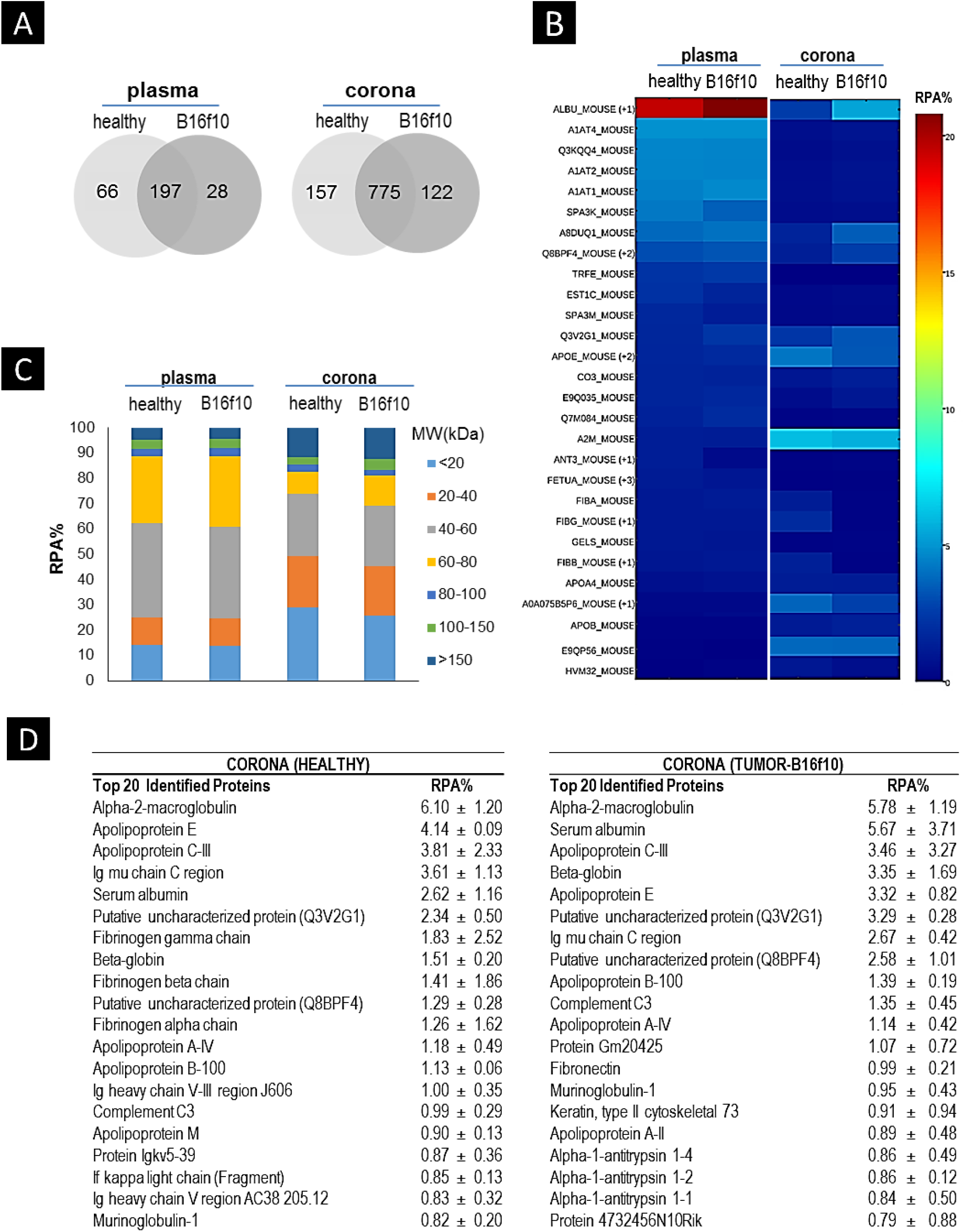
Comprehensive identification corona proteins associated with PEGylated liposomes in the blood circulation of healthy and melanoma-bearing C57 mice by LC-MS/MS. **(A)** Venn diagrams report the number of unique and common proteins between proteins identified in healthy and melanoma-bearing mice, in plasma control and corona samples. Peptide identifications were accepted if they could be established at greater than 50.0% probability and protein identifications were accepted if they could be established at greater than 99.0% probability and contained at least 2 identified peptides. Proteins shown in the Venn diagrams were identified in at least one of the three biological replicates; **(B)** Heatmap of Relative Protein Abundance (RPA %) values for plasma and corona proteins. Only proteins with RPA>1% in at least one of the samples are shown. Protein-rows are sorted according to the RPA% values (from highest to lowest) of the first sample (plasma healthy). The list of illustrated plasma and corona proteins and their respective accession numbers and RPA values are shown in Table S3; **(C)** Classification of plasma and corona proteins identified according to their molecular mass. The RPA% values for each molecular weight group represents the average of 3 biological replicates (n=3 mice/replicate); **(D)** Most abundant proteins (top-20) identified in the protein corona of liposomes intravenously injected in healthy and melanoma-bearing mice. RPA% values represent the average and standard deviation from 3 biological replicates (n=3 mice/replicate).

Of special interest for biomarker discovery is the low-abundance plasma proteome, secreted from the tumor microenvironment into the blood circulation, likely to contain the most clinically relevant and previously undiscovered markers. Only low MW intact proteins or protein fragments can passively diffuse through the endothelial barrier in the blood.^18^ However, the facile clearance of the low MW tumor-secreted proteins by kidney filtration limits their detection. We have previously shown that liposomes intravenously injected in healthy mice tend to interact with low MW plasma proteins.^11^ To examine if our previous finding applies also in the case of protein coronas extracted from tumor-bearing mice, the Relative Protein Abundance (RPA%) of each identified protein was calculated and proteins were then classified according to their molecular weight. As illustrated in Figure 2C, proteins with MW < 60 kDa accounted for approximately 70% of the protein coronas formed, in both healthy and tumor-inoculated mice. Remarkably, analysis of the *in vivo* protein coronas increased the identification of proteins with MW <40 kDa, in comparison with plasma control analysis (Figure 2C).

To explore the potential exploitation of the *in vivo* protein corona for biomarker discovery we compared protein coronas formed in healthy and melanoma-bearing mice, aiming to identify differentially abundant proteins. As revealed by the Venn diagram of Figure 2A, the composition of protein corona was different in the absence and presence of melanoma tumors, with 157 and 122 proteins uniquely found, respectively. Moreover, common proteins between the liposomal coronas formed in healthy and melanoma-bearing mice (n=775) i (Figure 2A), were not equally abundant, as depicted by the RPA (%) values of **Tables 2D** and **S4** and **Figure S1**. These results strengthened our hypothesis that proteomic analysis of the liposomal coronas reveals differences in healthy and diseased states and prompted us to further investigate if the above variations in the abundance of plasma proteins can be exploited for cancer diagnostics.

To assess the reproducibility of protein corona formation we compared the total number of identified proteins between the three biological replicates. As shown in **Figure S2**, ~60% and 55% of corona proteins were common between the three biological replicates in healthy and melanoma-bearing mice respectively. To identify those proteins that were reproducibly differentially abundant between healthy and melanoma-bearing mice in all the three biological replicates, further statistical analysis of the raw LC-MS/MS data was carried out using Progenesis software. The Relative Protein Expression (fold change) and the reliability of measured differences (ANOVA, *p value*) were calculated and are presented in Figure 3A and **Table S5**. Progenesis statistical analysis revealed 384 individual proteins differentially expressed (n=136 upregulated; n=248 downregulated) in tumor compared to healthy corona samples, with a *p value<0.05.* Astonishingly, the same comparative analysis performed for plasma samples, revealed only 6 potential biomarker proteins (**Table S6**). Overall, the above data suggest that proteomic analysis of the liposomal protein coronas eliminated the issue of albumin masking, increased the total number of plasma proteins identified and enabled the discovery of a significantly higher number of potential biomarkers, in comparison to plasma control analysis.

**Figure 3:**
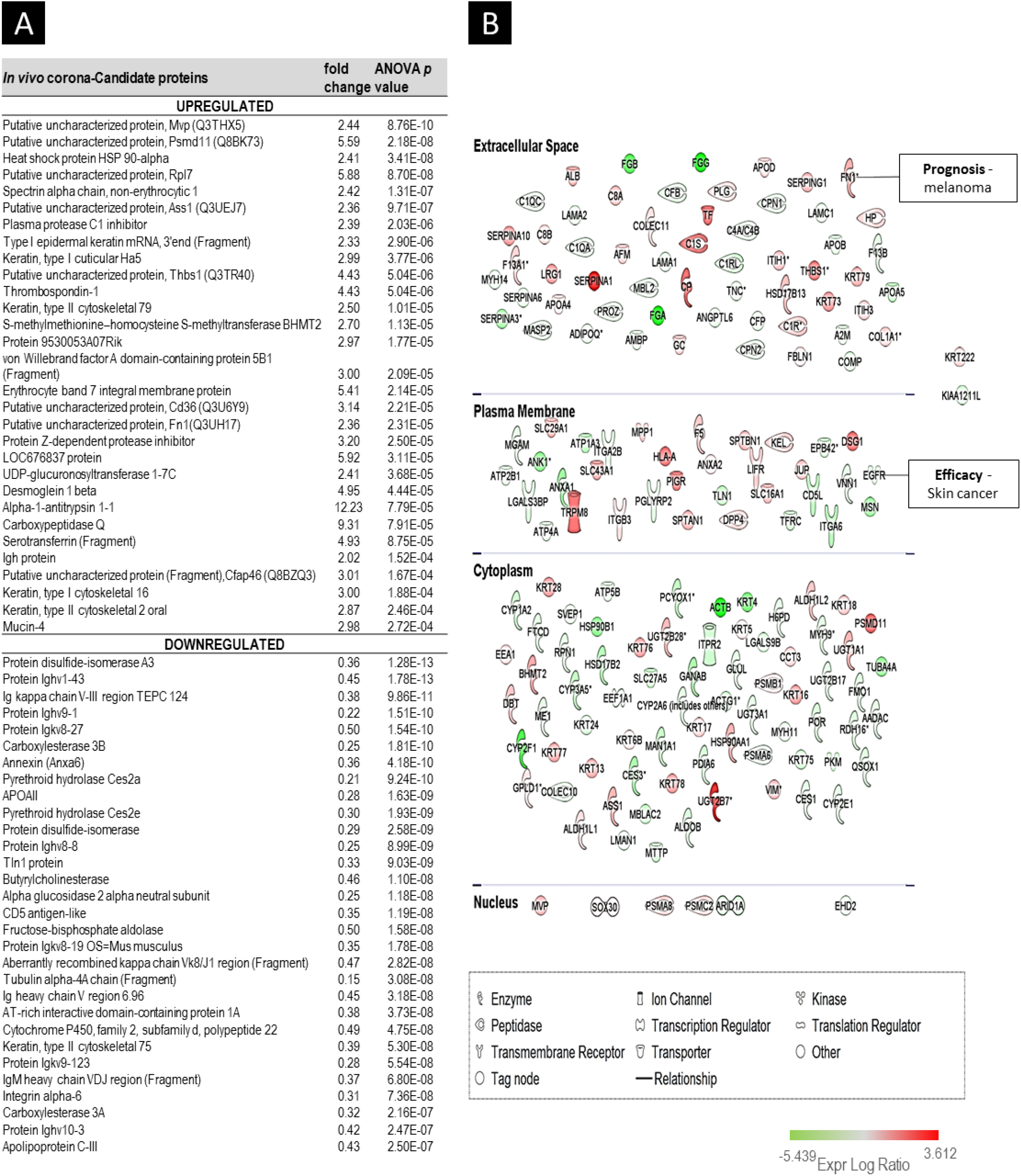
Candidate protein biomarkers, with differential abundance in the liposomal coronas formed in healthy and melanoma-bearing C57 mice. **(A)** Only proteins with at least 2 fold change with the lowest *p value* are shown). Data was filtered to a 1% false discovery rate (FDR). The full list of proteins with *p<0.05* are shown in **Table S5; (B)** Ingenuity Pathway Analysis (IPA) of potential biomarker corona proteins associated with melanoma cancer. Upregulated proteins (red; n=75) and downregulated proteins (green; n=95) are organised according to their cellular localisation (extracellular, plasma membrane, cytoplasmic, nuclear). The name of proteins illustrated in the diagram and their respective gene symbol, *p* values and fold change values are shown in Table S7.

To investigate whether the above 384 corona proteins have been previously associated with melanoma cancer, Ingenuity Pathway Analysis (IPA) was performed. Disease and function IPA search revealed the association of 170 different proteins with melanoma cancer pathways (with a *p value* of 1.24E-6) (Figure 3B and **Table S7**). According to IPA (Figure 3B), only 2 of them have been described in the literature as potential biomarkers for skin cancer (epidermal growth factor receptor and fibronectin 1). The ability of intravenously injected liposomes to surface-capture low abundance, disease specific molecules was further reinforced by the classification of the 170 melanoma-associated corona proteins according to their cellular localization. The detection of intracellular proteins, released from the tumor microenvironment into the blood, is a major challenge in proteomic analysis, due to their extremely low concentration in plasma.^19^ As shown in Figure 3B, 73 melanoma-associated proteins were found to be intracellular (n=71 cytoplasm; n=2 nucleus). Despite their involvement with previously described melanoma-associated cancer pathways, none of the intracellular corona proteins identified in this study has been reported in the literature as potential biomarker for melanoma diagnosis. Further evaluation will be necessary in order to define the sensitivity, specificity and predictive capabilities of these candidate biomarker proteins and to demonstrate their clinical utility.

### Characterization of the *in vivo* protein corona in lung carcinoma-bearing immune-deficient mice

Most cancers cause a wide range of host response mechanisms, resulting in the production of multiple inflammation-related proteins.^20^ Although these proteins could serve as an early index of tumorigenesis-induced inflammation, they lack of sufficient specificity and sensitivity for a single disease detection.^15^ Most of the high specificity cancer biomarkers are proteins shed from the tumor microenvironment cells into the blood circulation, however their detection in human plasma samples has proven to be extremely challenging due to their very low concentration in the ng/ml to pg/ml range. While syngeneic tumor models provide more consistent genetic and environmental backgrounds, inflammatory host responses might obscure the identification of tumor-released molecules.^21^ An alternative strategy that circumvents the host response problem is the inoculation of human tumors in immune deficient mice, which could allow the identification of tumor plasma markers derived from both cancer human cells and non-cancer host stromal cells simultaneously.^15-17^ The very low concentration of tumor released proteins in the blood, in addition to the albumin ‘masking’ effect have been the main challenges that compromise the use of xenograft models for biomarkers discovery.^22^

To assess the ability of intravenously administered liposomes to scavenge the blood pool and retrieve host-response mouse proteins as well as tumor released human proteins, we comprehensively compared protein coronas formed around liposomes (10 min post-injection) in healthy and lung carcinoma-bearing SCID mice (4 weeks post-inoculation of the A549-luc cells). Protein corona fingerprints characterised for healthy and diseased mice were found to differ not only quantitatively (Figure 1C) but also qualitatively, as depicted in Figures 4A and 4B. As shown in the Venn diagram of Figure 4A, 105 and 85 proteins were uniquely found in the absence and presence of lung carcinomas, respectively. Moreover, corona proteins commonly identified in healthy and lung-carcinoma mice (n=847) were not equally abundant (Figure 4B **and Table S8**). In agreement with our previous results, protein coronas formed in SCID mice were mainly composed of low MW proteins (>60% of corona proteins had a MW<60 kDa); (Figure 4C). The above data, are in agreement with the results obtained for the melanoma model and demonstrate the potential use of this technology for biomarker discovery in the presence and absence of a functional immune system.

**Figure 4:**
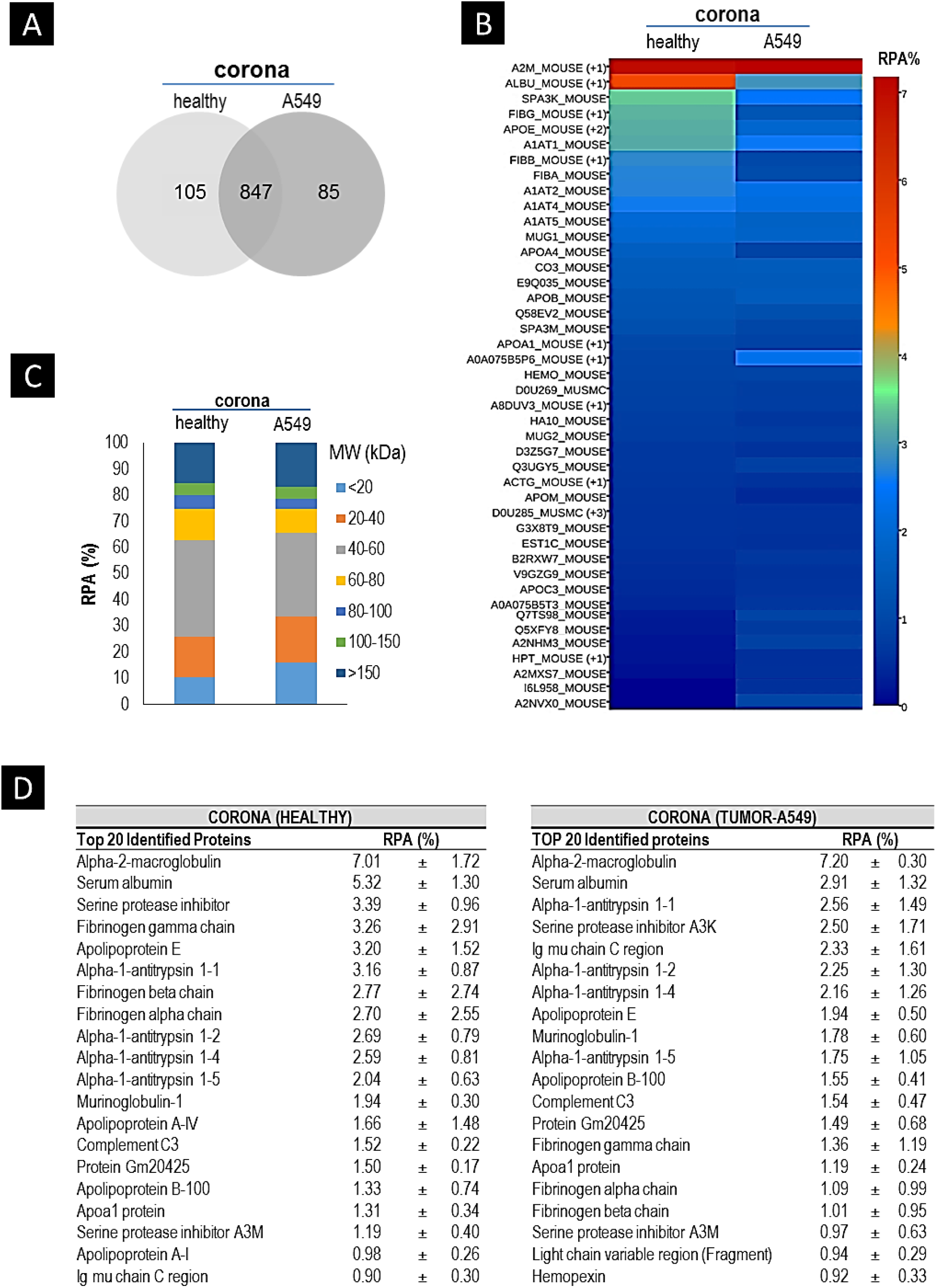
Comprehensive identification of corona proteins associated with liposomes in the blood circulation of healthy and lung carcinoma-bearing SCID nude mice. **(A)** Venn diagrams report the number of unique and common proteins between proteins identified in the coronas formed onto liposomes intravenously injected in healthy and lung-carcinoma bearing mice. Peptide identifications were accepted if they could be established at greater than 50.0% probability and protein identifications were accepted if they could be established at greater than 99.0% probability and contained at least 2 identified peptides. Proteins shown in the Venn diagrams were identified in at least one of the three biological replicates; **(B)** Classification of plasma and corona proteins identified according to their molecular mass. The percentage of relative protein abundance (%RPA) for each molecular weight group represent the average of 3 biological replicates (n=3 mice/replicate); **(C)** Heatmap of RPA (%) of proteins identified in the coronas formed onto liposomes intravenously infused in healthy and lung-carcinoma mice, as identified by LC-MS/MS. Only proteins with RPA>0.5 % are shown. RPA (%) values represent the average of 3 biological replicates (n=3 mice/replicate). The full list of corona proteins identified and their respective accession numbers are shown in **Table S8**; **(D)** Most abundant proteins (top-20) identified in the protein corona of liposomes intravenously injected in healthy and lung carcinoma-bearing mice. Relative protein abundance (RPA) values represent the average and standard deviation from 3 biological replicates (n=3 mice/replicate).

Progenesis statistical analysis of the corona proteins revealed 328 proteins differentially expressed with statistical significance (*p value* <0.05) between tumor-bearing and healthy animals, of which 178 were upregulated and 150 were downregulated (Figure 5A and **Table S9**). In addition, disease and function IPA search demonstrated the association of 172 different proteins with previously reported adenocarcinoma pathways (with a *p value* of 2.6E-5); (**Figure S3**), and the association of 43 proteins with lung cancer pathways, as shown in Figure 5B. According to IPA, 8 out of the 43 lung cancer related corona proteins have been already described in the literature as potential biomarkers (3 for diagnosis, 1 for treatment efficacy, 1 disease progression, 1 safety and 1 unspecified application) (Figure 5B). It is becoming increasingly evident that tumor progression is determined by a number of complex interactions between malignant cells and their surrounding endogenous host stromal cells.^23^ The identification of intracellular and lung-carcinoma associated proteins (Figure 5B), suggests that the nanoscale liposomal corona contains host (murine) stromal released proteins.

**Figure 5:**
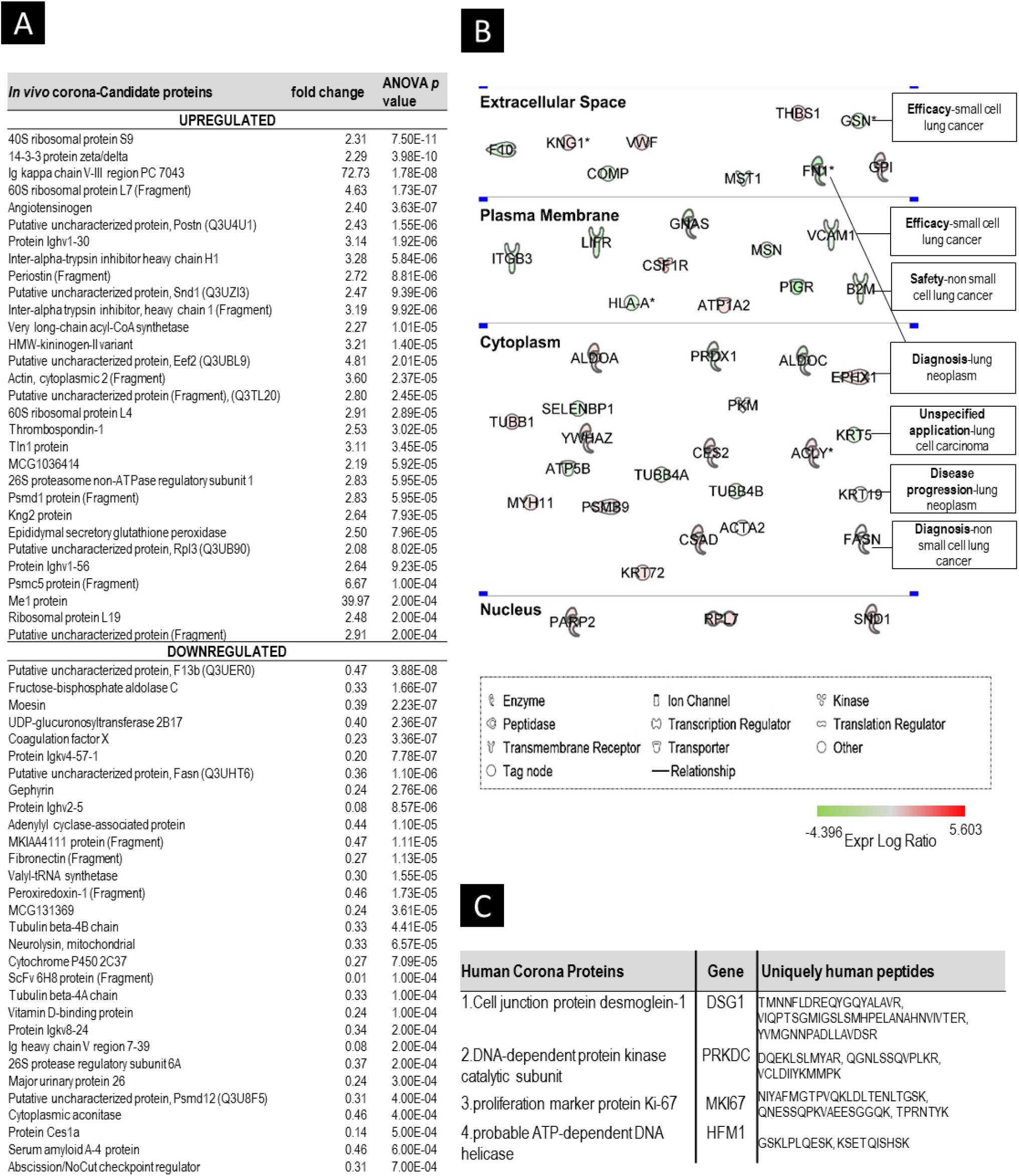
Candidate protein biomarkers, with differential abundance in the liposomal coronas formed in healthy and lung carcinoma-bearing SCID nude mice. **(A)** Only proteins with at least 2 fold change with the lowest *p value* are shown. The full list of proteins with *p<0.05* are shown in Table S9. Data was filtered to a 1% false discovery rate (FDR); **(B)** Ingenuity Pathway Analysis (IPA) of potential biomarker corona proteins associated with lung carcinoma. Upregulated proteins (red; n=21) and downregulated proteins (green; n=22) are organised according to their cellular localisation (extracellular, plasma membrane, cytoplasmic, nuclear). The name of proteins illustrated in the diagram and their respective gene symbol, *p* values and fold change values are shown in Table S10; **(C)** List with human proteins (secreted from A549-luc cells) identified in the coronas of intravenously injected liposomes in lung-carcinoma bearing mice. Only proteins with two or more human-specific peptides are shown.

Finally, we wanted to investigate whether any uniquely human proteins, released from the transplanted A549 cells into the blood circulation of SCID mice, were found in the liposomal corona. All corona proteins identified, were sorted into ‘human’, ‘mouse’, or ‘indistinguishable’ based on their species-specific sequences. To confirm the species assignment, human peptides were searched using BLAST against a mouse-only database (UniProt/SwissProt) to remove any human sequences that exactly matched mouse sequences. When considered the detection of human-sourced protein material to be evidenced by the detection of two or more human specific peptides per protein identity (Figure 5C), four human proteins were found to be surface-captured by liposomes. Among them, proliferation marker protein Ki-67 was previously described as a potent biomarker with significant predictive and prognostic value in lung cancer.^24^ The identification of human proteins in the mouse serum of xenograft models has been scarcely attempted previously and required the extensive multidimensional fractionation of serum proteins prior to proteomic analysis.^15^ In this study, our results suggest that liposomes can act as carriers not only for low abundance blood circulating proteins released from the host cells of the tumor microenvironment, but also for highly specific biomarker proteins shed by tumor cells.

## Discussion

New molecular biomarkers are needed to improve cancer diagnosis and evaluate disease progression and response to treatment.^25^ The proteomic technologies available today are powerful tools for cancer biomarker detection. However, the discovery of blood biomarkers using proteomics is hampered by the wide dynamic range of blood proteins which spans more than ten orders of magnitude, with tumor-tissue derived proteins present at the lower end of this range.^2^ Robust high-throughput proteomic platforms to facilitate the identification of blood-buried molecules are of immediate importance. Nanotechnology-enabled discovery of novel biomarkers, holds great promise but it is still in its infancy. Previously described nanoparticle-based technologies have been mainly used to detect already known disease-specific molecules.^26-29^ Even though NPs have been found to spontaneously interact with a wide range of different protein molecules once in contact with biofluids, the surface-capture of disease-specific molecules by blood circulating NPs has never been attempted before.^6^

The concept of utilizing the nanoparticle protein corona fingerprints for biomarker discovery has been only theoretically proposed and remains experimentally unexplored,^30^ with the exception of a recent report by Miotto *et al.,* who investigated protein corona formation after the *ex vivo* incubation of active maghemite nanoparticles (SMANS) in the milk of cows affected by mastitis.^31^ Whether the NPs employed in this study are able to harvest disease specific molecules from more complex and clinically-relevant biofluids, such as plasma, remains to be explored. The idea of utilizing protein corona for biomarker discovery was further reinforced by preliminary *ex vivo* studies suggesting that changes in the plasma proteome, induced by different disease phenotypes, may be reflected in the protein corona.^32, 33^ However, no report today has provided comprehensive MS-based identification of the proteins that consists such ‘personalized’ NP coronas.

In this study, we aimed to offer a comprehensive comparison between the protein coronas that formed *in vivo* onto clinically used liposomes in healthy and tumor-bearing mice, by LC-MS/MS. Our results, provided the first evidence that protein corona fingerprinting indeed varies, both quantitatively (Figure 1C) and qualitatively (Figures 2B and 4B), in the absence and presence of tumor growth. These differences observed between the protein coronas formed in melanoma and human lung carcinoma models compared to healthy control mice, supported the hypothesis that the interaction of liposomes with blood-circulating proteins is greatly affected by the ongoing pathophysiological events.

High abundance, high MW proteins, such as albumin and immunoglobulins present in the blood circulation hinder the detection of the low MW blood proteomic fractions, likely to contain previously unidentified disease-specific biomarkers. In addition, the rapid clearance of small proteins from the blood circulation by kidney filtration is another reason why the low MW region of the blood proteome remains largely unexplored. The only way a small molecule can remain in the blood circulation for longer periods is to adhere to a long-circulating, high abundance protein, such as albumin. Plasma depletion methodologies, often used as a prefractionation tool to address the issue of the ‘large dynamic concentration range’ of plasma proteome, discard those carrier proteins and inevitably their valuable cargos.^34^

Our data in this study suggest that the liposomal corona results in an ‘enriched’ sampling of the blood proteome, by minimising the ‘noise’ from highly abundant proteins, contrary to plasma control (Figure 1D). It should be emphasized that albumin was not depleted from plasma control or corona samples. Proteomic analysis of the recovered *in vivo* corona resulted in a significantly increased number of identified proteins in comparison to conventional proteomic analysis of plasma samples (Figure 2A). Even though albumin and other highly abundant plasma proteins were found to interact with the surface of liposomes (in both healthy and tumor-bearing mice) the extensive purification processes applied to retrieve the corona-coated liposomes and purify them from the unbound proteins, worked as fractionation tool and increased the range of plasma protein detection (Figures 2A and 2B).

Interestingly, classification of the corona proteins according to their molecular weight demonstrated the high affinity of the liposomal surface for interaction with low MW proteins (Figures 2C and 4C). It is possible that the low MW proteins identified have high affinity and interact directly with the surface of PEGylated liposomes (also known as ‘hard’ corona proteins) and/or they are trapped between other corona-carrier proteins that are adhered to the NPs surface (also known as ‘soft’ corona proteins).^35^ It should be emphasized, that the manner in which proteins adsorb and self-assemble onto the NPs surface is highly dependent on the NPs physicochemical properties including their size, surface curvature and functionalization. Albeit the most crucial determinant factors have been defined, concrete relationships between nanomaterials synthetic identity and protein corona composition in complex physiological environments remain elusive.^6^ Distinct proteins could be either enriched or displayed only weak affinity for the NP surface depending on the balance between the rates of association (Kon) and dissociation (Koff). Whether different nanomaterials could allow the detection of distinct biomarker signatures remains to be investigated in future studies.

Given the substantial heterogeneity among cancers, it is unlikely that any single protein molecule will have adequate specificity and sensitivity needed for accurate early diagnosis. There is a growing consensus that multiple markers, used individually or as part of a panel, are required for early disease detection. To better comprehend the potential use of the *in vivo* protein corona for biomarker discovery, we statistically compared the level of proteins present in the coronas formed in healthy and tumor-bearing mice. Progenesis analysis revealed 384 and 328 different proteins differentially expressed in tumor compared to healthy corona samples, in the melanoma (Figure 3A) and lung-carcinoma model, respectively (Figure 5A). It should be noted that the equivalent analysis of plasma samples in the case of melanoma model, revealed only 6 potential biomarker proteins (**Table S6**).

Proteins released from the tumor microenvironment, by leakage or secretion, are massively diluted in the blood circulation and current proteomic methodologies often fail to provide sufficient depth of analysis, required for their detection. Classification of the melanoma and lung carcinoma associated corona proteins identified in this study according to their cellular localization, demonstrated that circulating liposomes act as platforms for interaction with intracellular, tissue leakage products. This was further exemplified by the detection of human corona proteins released by lung carcinoma cells in the blood circulation of SCID mice (Figure 5C).

On a broader context, our findings suggest that the adsorption of small biomolecules onto the surface of NPs, once injected in the bloodstream allows thus their successful detection. The nanoscale liposomal corona is enriched by multiple plasma proteins with potential diagnostic and disease monitoring value which renders the need for much more work on the biomarker verification and validation front necessary, but beyond the scope of this study. The technology proposed here facilitates the initial discovery phase of the cancer biomarker pipeline in which animal models are mainly used, aiming to reveal molecules differentially abundant between healthy and diseased states. When animal models are employed for plasma biomarker discovery, the exploitation of the molecularly richer *in vivo* protein corona is beneficial in comparison to its counterpart *ex vivo* corona, however further investigations are required to assess the potential use of the *ex vivo* corona directly from human plasma samples. It is also important to understand the potential limitations of characterising protein corona fingerprints at a single time point post-inoculation of tumor cells in mice and more studies are needed in order to investigate the evolution of protein corona at different stages of tumor growth and prove its potential at the early asymptomatic stages of cancer.

## Conclusion

In this study, we provide previously unreported experimental evidence of the potential exploitation of the spontaneous and often undesirable protein coating of NPs, once injected in the blood circulation, for biomarker discovery. We demonstrated the ability of intravenously administered clinically used NPs to scavenge the blood pool of melanoma (B16-F10) and lung carcinoma-bearing (A549) mice and surface-capture a complex mixture of tumor-specific proteins that could not be identified by plasma sample analysis that was performed in comparison. The successful recovery and purification of corona-coated liposomes from the blood circulation of mice allowed the elimination of the overwhelming signal of highly abundant plasma proteins and enabled the detection of low abundant and low MW tumor-released proteins. Protein corona was found to qualitatively and quantitatively differ in the absence and presence of inoculated tumors enabling the discovery of 384 and 328 potential biomarkers for melanoma and lung adenocarcinoma respectively. The success of the nanoparticle-enabled retrieval of low abundance plasma proteins was further revealed by the concurrent detection of murine host response proteins and human inoculated tumor-released proteins in a lung adenocarcinoma xenograft model.

## Experimental

### Preparation of liposomes

Two different batches of HSPC:Chol:DSPE-PEG2000 (56.3:38.2:5.5) liposomes were prepared, one for the melanoma and one for the lung adenocarcinoma model by thin lipid film hydration method followed by extrusion, as we have previously described.^10, 11^ The same batch of liposomes was used for all the three biological replicates of each tumor model. The physicochemical properties of liposomes are shown in **Tables S1** & **S2**.

### Cell culture

B16f10-luc, a murine melanoma cancer cell line and A549-luc, a human lung carcinoma cell line were bought from Caliper Life Sciences, Inc. Cells were cultured in Advanced RPMI (GIBCO) supplemented with 10% FBS (Fetal Bovine Serum) and 1% glutamine. Cells were grown in a humidified 37 °C/5% CO2 incubator and passaged at 80% confluence.

### Animals and tumor models

Six to eight week old male nude SCID beige and five to six week old female C57BL6 were purchased (Charles River, UK) and acclimatized to the environment for at least 7 days before any experimental procedures. Animal procedures were performed in compliance with the UK Home Office Code of Practice for the Housing and Care of Animals used in Scientific Procedures.

For each tumor model three biological replicates were performed, each using 3 mice (n=3 biological replicates, n=3 mice/replicate). The plasma samples recovered from three mice were pooled for each biological replicate). Healthy SCID and C57BL6 mice were also used as a control (n=3 biological replicates, n=3 mice/replicate). The melanoma tumor model was established by subcutaneous injection of 1 × 10^7^ B16F10-luc cells in a volume of 50 μl of PBS into the left leg of C57BL6 mice. The lung carcinoma model was established by the intravenous injection (via tail vein) of 5 × 10^6^ A549-luc cells in a volume of 200ul, in SCID mice. The same batch of cells was prepared and injected for all three biological replicate of each tumor model.

For both models tumor growth was monitored by IVIS Lumina II imaging system (Caliper Life Sciences Corp., Alameda, CA). Luciferin (150 mg/kg in PBS; VivoGlo™, Promega) was intraperitoneally injected 12 minutes before imaging as a substrate for the luciferase expressing tumour cell lines. In the case of lung adenocarcinoma model liposomes injections were performed when measurable bioluminescence signal was detected in the lungs of all mice. For the melanomal model, the tumour volume was also estimated by measuring three orthogonal diameters (a, b, and c) with calipers and calculated as (a×b×c) × 0.5 mm3. Liposome injections were performed at a tumour volume of 200-400 mm^3^.

### Protein corona formation after *in vivo* administration

Liposomes were administered intravenously via the lateral tail vein (at a lipid dose of 0.125mM/g body weight) in healthy control and tumor-bearing mice after anaesthesia with isoflurane. Ten minutes post-injection, blood containing corona-coated liposomes was recovered by cardiac puncture in K_2_EDTA coated collection tubes. ~0.5 ml of blood sample was collected from each mouse. Plasma was then prepared by centrifugation for 12 minutes at 1200 RCF at 4 °C after inverting the collection tubes to ensure mixing of blood with EDTA. Plasma was the collected into Protein LoBind Eppendorf Tubes. The plasma samples obtained from three mice were pooled together for a final plasma volume of 1 ml.

### Separation of corona-coated liposomes from unbound and weakly bound proteins

Liposomes recovered from plasma were separated form excess plasma proteins by size exclusion chromatography followed by membrane ultrafiltration as we have previously described.^10-12^

### Size and zeta potential measurements using dynamic light scattering (DLS)

Liposome size and surface charge were measured using Zetasizer Nano ZS (Malvern, Instruments, UK).

### Transmission Electron Microscopy (TEM)

Liposomes were visualized with transmission electron microscopy (FEI Tecnai 12 BioTwin) before and after corona formation using Carbon Film Mesh Copper Grid (CF400-Cu, Electron Microscopy Science), after staining with aqueous uranyl acetate solution 1%.^10^

### SDS-PAGE electrophoresis

Corona proteins associated with 0.025 μM of liposomes and plasma samples (5 ul) were loaded and run in a 4-20% Novex® Tris-Glycine SDS Running Buffer (ThermoFisher Scientific), according to manufacturer’s instructions. The gels were stained with Imperial Gel Staining reagent (Sigma Life Science).

### Quantification of adsorbed proteins

Proteins associated with recovered liposomes were quantified by BCA Protein assay kit and the amount of liposomes recovered by Stewart assay.^10-12^ Pb values, expressed as μg of protein/μmole of lipid were calculated and presented as the average ± standard deviation (n=3 biological replicates, n=3 mice/replicate). For the comparison of Pb values (Figure 1C) statistical analysis of the data was performed using GraphPad Prism 7.02 software. Unpaired two-tailed t-test was performed and p values < 0.05 were considered significant.

### Mass Spectrometry

In-gel digestion of corona (40ug) and plasma (5ul) proteins was performed prior to LC-MS/MS analysis, as we have previously described.^11-13^ UltiMate® 3000 Rapid Separation LC (RSLC, Dionex Corporation, Sunnyvale, CA) was coupled to an Orbitrap Fusion™ (Thermo Fisher Scientific, Waltham, MA) mass spectrometer.

### Mass Spectrometry data analysis

Data produced were searched using Mascot (Matrix Science UK), against the [SwissProt_2016_04 database]. The Scaffold software (version 4.3.2, Proteome Software Inc.) was used to validate MS/MS based peptide and protein identifications and for relative quantification based on spectral counting (Figures 2 & 4). Semi quantitative assessment of the protein amounts was conducted using normalized spectral counting (50% peptide probability, 99% protein probability, at least 2 identified proteins), as we have previously reported.^10-13^ Heatmaps of RPA values were prepared using Plotly 2.0 software.

To statistically compare proteins identified in the coronas formed in healthy and tumor-bearing mice (Figures 3 & 5), MS peak intensities were analyzed using Progenesis LC-MS software (version 3.0; Nonlinear Dynamics). Data was filtered to a 1% false discovery rate (FDR). The peptide intensities were compared between groups by one way analysis of variance (ANOVA, p<0.05 for statistical significance).

Mass Spectrometry data were analysed through the use of QIAGEN’s Ingenuity® Pathway Analysis (IPA®, QIAGEN Redwood City, www.qiagen.com/ingenuity). Diseases and functions IPA tool was used to identify proteins involved in melanoma and lung-carcinoma associated proteins. The biomarker overlay IPA tool was then used to identify proteins described in the literature as potential biomarkers for melanoma and lung cancer.

### Species-specific peptide identification

Mass spectrometry datasets were analysed using Mascot and Scaffold as described above, and peptide lists were generated with either mouse or human taxonomy selected. Peptides were classified as being unique to mouse or human databases, or common to both using in-house code (Mathematica version 11.0.1; Wolfram Research, Champaign, IL).^36^ Peptide species were checked using the BLAST protein database (National Center for Biotechnology Information, Bethesda, MD). We considered the detection of human-sourced protein material to be evidenced by the detection of two-or-more human-specific peptides per protein identity. Peptides were aggregated over triplicate samples; any peptides that were also detected in triplicate mouse-only control samples (i.e. false positives) were removed from consideration; peptides differing between mouse and human by only a single amino acid substitution were also not considered.

## Author contributions

M.H. initiated, designed and performed all the experiments, analysed all data obtained and wrote the manuscript. Z.A aided in the injections of tumor cells and IVIS imaging. M.B performed the cell culture experiments. J.S aided in the species-specific peptide identification. K. K. initiated, and directed this study, provided intellectual input and contributed to the writing of the manuscript.

## Acknowledgements

This study was partially funded by the Marie Curie Initial Training Network PathChooser (PITN-GA-2013-608373). Authors would like to thank the Faculty of Life Sciences EM Facility at the University of Manchester. and Leo Zeef for his help with IPA analysis.

## Supporting Information Available

Table S1: The effect of protein corona formation on the physicochemical characteristics of liposomes (melanoma model); Table S2: The effect of protein corona formation on the physicochemical characteristics of liposomes (lung adenocarcinoma model); Table S3: Blood-circulation proteome analysis. List of proteins identified in plasma and corona samples recovered from healthy and melanoma-bearing C57 mice with RPA>1% in at least one of the samples; Figure S1: Comparison of protein coronas formed in healthy and melanoma bearing C57 mice, as identified by LC-MS/MS; Table S4: Comparison of protein coronas formed in healthy and melanoma-bearing C57 mice; Figure S2: Reproducibility of protein corona formation; Table S5: Candidate melanoma biomarkers, as identified by proteomic analysis of the liposomal coronas; Table S6: Candidate melanoma biomarkers, as identified by proteomic analysis of plasma samples; Table S7: Corona proteins-potential biomarkers associated with melanoma cancer, according to Ingenuity Pathway Analysis (IPA); Tables S8: Comparison of protein coronas formed in healthy and lung cancer-bearing SCID mice; Table S9: Candidate lung carcinoma biomarkers, as identified by proteomic analysis of the *in vivo* liposome coronas; Table S10: Corona proteins-potential biomarkers associated with lung carcinoma, according to Ingenuity Pathway Analysis (IPA); Figure S3: Ingenuity Pathway Analysis (IPA) of proteins associated with adenocarcinoma identified in the liposomal corona; Table S11: Corona proteins-potential biomarkers associated with adenocarcinoma, according to Ingenuity Pathway Analysis (IPA).

